# Systematic review of *Plasmodium falciparum* and *Plasmodium vivax* polyclonal infections: Impact of prevalence, study population characteristics, and laboratory procedures

**DOI:** 10.1101/2021.03.18.436023

**Authors:** Luis Lopez, Cristian Koepfli

## Abstract

Multiple infections of genetically distinct clones of the same *Plasmodium* species are common in many malaria endemic settings. Mean multiplicity of infection (MOI) and the proportion of polyclonal infections are often reported as surrogate marker of transmission intensity, yet the relationship with traditional measures such as parasite prevalence is not well understood. We have searched Pubmed for articles on *P. falciparum* and *P. vivax* multiplicity, and compared the proportion of polyclonal infections and mean MOI to population prevalence. The impact of the genotyping method, number of genotyping markers, method for diagnosis (microscopy/RDT vs. PCR), presence of clinical symptoms, age, geographic region, and year of sample collection on multiplicity indices were assessed. For *P. falciparum*, 153 studies met inclusion criteria, yielding 275 individual data points and 33526 genotyped individuals. The proportion of polyclonal infections ranged from 0-96%, and mean MOI from 1-6.1. For *P. vivax*, 54 studies met inclusion criteria, yielding 115 data points and 13325 genotyped individuals. The proportion of polyclonal infections ranged from 0-100%, and mean MOI from 1-3.8. For both species, the proportion of polyclonal infections ranged from very low to close to 100% at low prevalence, while at high prevalence it was always high. Each percentage point increase in prevalence resulted in a 0.34% increase in the proportion of polyclonal *P. falciparum* infections (*P*<0.001), and a 0.78% increase in the proportion of polyclonal *P. vivax* infections (*P*<0.001). In multivariable analysis, higher prevalence, typing multiple markers, diagnosis of infections by PCR, and sampling in Africa were found to result in a higher proportion of *P. falciparum* polyclonal infections. For *P. vivax*, prevalence, year of study, typing multiple markers, and geographic region were significant predictors. In conclusion, polyclonal infections are frequently present in all settings, but the association between multiplicity and prevalence is weak.

## Introduction

Malaria remains a global public health threat with over 200 million confirmed cases and over 400,000 deaths recorded in 2019 [1]. Reliable measurements of transmission intensity are needed in order to optimally allocate resources for control and to measure changes response to control. Obtaining such measures is a challenge [2].

Individuals in malaria endemic regions are often concurrently infected with several clones or strains of the same parasite species. Polyclonal infections can be the result of a single mosquito bite transmitting a genetically diverse inoculum (coinfection), or of repeated bites transmitting different clones (superinfection) [3]. In *P. vivax*, relapses of heterologous hypnozoites during ongoing blood-stage infection can also result in polyclonal infections [4, 5].

Multiplicity of infection (MOI, also termed ‘complexity of infection’, COI), and the proportion of polyclonal infections (also termed ‘multiple clone infections’) are among the most frequently reported measures of molecular malaria surveillance studies of *Plasmodium falciparum* and *Plasmodium vivax*. A large number of genotyping protocols is used to determine multiplicity. The most common methods include typing of size-polymorphic markers (antigens or microsatellites) and sizing on agarose gel or by capillary electrophoresis (CE) [6, 7], SNP panels [8], and more recently amplicon sequencing [9, 10].

MOI is generally assumed to be higher in high-transmission settings, while in low-transmission settings most individuals might carry single clone infections. Multiplicity has thus been proposed as surrogate marker to measure changes in transmission intensity in time and space. Numerous studies found a correlation between transmission intensity and MOI. For example, in West Papua, Indonesia, clinical incidence of *P. vivax* and *P. falciparum* approximately halved over a 14-year period. The proportion of polyclonal infections fell from 70% to 35% for *P. vivax*, and from 28 to 18% for *P. falciparum* [11]. Along the Thai-Myanmar border, a 10-fold decrease in incidence over a 10-year period was accompanied by a reduction in the proportion of polyclonal *P. falciparum* infections from 63% to 14% [12]. In Senegal, the proportion of polyclonal infections fell from 50% in 2001–2002 to 0% in 2014 along with a decrease in incidence [13]. In Venezuela, multiplicity increased dramatically in parallel to an increase in incidence [14]. In Burkina Faso, MOI mirrored seasonal patterns of incidence [15]. In other studies, however, the association between multiplicity of infection and transmission intensity was weak [16], or absent [17].

Understanding the extent of polyclonal infections is important for molecular malaria surveillance activities beyond measuring transmission intensity. Typing of molecular markers for drug resistance is an important aspect of malaria surveillance [18]. Polyclonal infections can result in estimates of prevalence (i.e. the proportion of all individuals carrying a drug resistant parasite) to differ substantially from the true frequency (i.e. the proportion of all parasite strains in the population carrying the marker of resistance) [19]. Rapid diagnostic tests (RDTs) are key tools for malaria diagnosis. Most RDTs for *P. falciparum* diagnose the HRP2 protein. Increasing numbers of parasites are being reported carrying a deletion of the *hrp2* gene, which results in false negative tests. Molecular surveillance of *hrp2* deletion is a high priority [20]. In case of polyclonal infections, wild type parasites can mask strains carrying the deletion, resulting in an underestimation of the frequency of deletion [21]. Changes in the frequency of polyclonal infections could result in observed changes in the prevalence of deletions and mutation, even if population frequencies remain constant. Data on multiplicity is also required for the reconstruction of haplotypes from whole genome sequencing data. In the presence of polyclonal infections, specific software tools are required [22]. As a further line of research, the impact of polyclonal infection on clinical symptoms has been studied [23-25].

Despite the large number of studies reporting multiplicity data, the relationship between MOI or the proportion of polyclonal infections and transmission intensity across a wide range of studies remains poorly characterized. Likewise, little effort has been made to systematically assess the impact on multiplicity estimates of characteristics of the study population (e.g. asymptomatic vs. clinical infections) or the method used for genotyping (e.g. gel electrophoresis vs. capillary electrophoresis for sizing of polymorphic markers). We have conducted a systematic literature search and compared the proportion of polyclonal infections and mean multiplicity to population prevalence of infection as common measure of transmission intensity. Furthermore, we studied the effect of laboratory procedures and characteristics of the study population on multiplicity indices.

## Methods

### Search Strategy

PubMed was searched on 1/10/21 using the searches “Multiplicity of infection *Plasmodium falciparum*”, “Complexity of infection *Plasmodium falciparum*”, and “Multiple clone infection *Plasmodium falciparum*”. Articles were eligible if they reported data on mean MOI and/or the proportion of polyclonal infections. From clinical trials, data was included only when reported separately for the baseline. 265 studies were identified as eligible for assessment out of which 153 were included in the pooled analysis.

The same search strategy for *P. vivax* yielded very few results. Thus, a broader search attempt was made. This included screening of additional studies from authors that were identified in the initial search for *P. falciparum* or *P. vivax* data, and screening of studies that cited widely used methods for *P. vivax* genotyping (e.g. [7, 26, 27]). The final dataset for *P*. vivax included 54 studies.

From each study, data on multiplicity of infection and/or the percentage of polyclonal infections were recorded. If available, population prevalence of infection was recorded from the article. If no prevalence data was reported, data was obtained from the Malaria Atlas Project (MAP) website (https://malariaatlas.org). MAP provides modeled prevalence of infection by microscopy in 1-10-year-old children at the country or subcountry level. For the current study, data from the lowest administrative level available was recorded. With very few exceptions data points for with prevalence data was taken from MAP had only genotyped children. Thus, the age range of the individuals genotyped matched the age range of the data in MAP. For studies conducted before the year 2000, data from 2000 was recorded (the earliest available from MAP).

Additional elements of the eligible articles that were analyzed included the country of study, the year of sample collection, whether the study population was clinical or asymptomatic, the method for diagnosis, the method for genotyping, and the age range of the study population. The total number of patients surveyed, and the number of patients genotyped was recorded. A few studies did not report genotyping success rate, and thus the total number of genotyped samples is unknown. In these cases, it was assumed that all infections underwent genotyping, and the number of samples with *P. falciparum* and *P. vivax* infections was included.

Certain articles that met the inclusion criteria yielded more than one data point. Elements within studies that prompted individual treatment included examination of multiple geographic locations, investigation of both clinical and asymptomatic patients separately, inclusion of multiple age ranges, or reporting data from multiple different time periods.

### Data Analysis

In addition to prevalence of infection, characteristics of the study population and the laboratory methods selected for diagnosis and genotyping can affect MOI and the proportion of polyclonal infections. The impact of the following factors on multiplicity indices was analyzed:

(i) Number of genotyping markers: A larger number of molecular markers increases the probability to detect additional clones. Studies were distinguished between using a single marker or multiple markers. SNP panels always include multiple markers, but only two alleles can be distinguished for each SNP. Whole genome sequencing (WGS) covers the entire genome, but read depth is generally lower than amplicon sequencing. Thus, SNP panels and WGS were excluded from this analysis. (ii) Diagnosis: In all transmission settings, a large proportion of infections remains below the limit of detection of microscopy or rapid diagnostic test (RDT) and can only be diagnosed by sensitive molecular methods (e.g. PCR) [28, 29]. Detectability of minority clones is higher when parasite density is higher [30]. While individual data on parasite density was not available, data was grouped into infections diagnosed either by PCR (i.e. more low-density infections included) or diagnosed by microscopy or RDT. (iii) Genotyping method: Estimates of multiplicity are heavily impacted by the ability to detect minority clones. Capillary electrophoresis generally increases detectability of minority clones and distinction of fragments of similar size compared to visualization on gel, thus CE detects higher multiplicity [31, 32]. Amplicon deep sequencing further increases the ability to discriminate between clones [9]. Data was thus grouped by genotyping method as follows: a) size polymorphic markers visualized on gel, b) size polymorphic markers visualized by CE, c) SNP panels, d) amplicon sequencing, and e) whole genome sequencing. (iv) Clinical vs. asymptomatic infections: Parasite density in clinical samples is expected to be higher than in asymptomatic samples, affecting the ability to detect minority clones. Thus, the effect of sampling from clinical vs. asymptomatic populations was assessed. (v) Age: Multiplicity of infection has been shown to be affected by age [33]. For the analysis, data was grouped into age ranges as follows: including individuals of all ages, only individuals ≤15 years, and only individuals >15 years included. Some studies that did not fully fit this classification were assigned to the closest matching group (supplementary file S1). In moderate and high transmission settings, clinical disease occurs mostly in children, in particular in the case of *P. vivax*. Thus, it is expected that many studies on clinical patients enrolled mostly children. Nevertheless, unless an age limit was stated, they were classified as including all ages. (vi) Geographic locations: Studies were grouped as conducted either in Africa, the Asia-Pacific, or the Americas. (vii) Year: Studies conducted from 1990 to 2019 were included in the analysis. Over this period, the number of *P. falciparum* and *P. vivax* cases globally has decreased [1]. Associations between the year of sample collection and multiplicity of infection indices were analyzed. If the year of sample collection was not mentioned, the year prior to the year of publication was used for analysis.

Linear regression models were used to analyze the impact of the above-described variables and the proportion of polyclonal infections or mean multiplicity. Multivariable regression was conducted, with step-wise removal of variables with a *P*-value >0.05, until the final model included only variables with *P*≤0.05.

## Results

### *P. falciparum* literature search

The literature search yielded 153 studies that met the inclusion criteria, resulting in 275 data points with a total of 33,526 genotyped infections. Summary statistics are presented in Table 1. On average, 122 individuals were genotyped per data point. The proportion of polyclonal infections ranged from 0-99%, and mean MOI from 1-6.1.

**Table 1:**
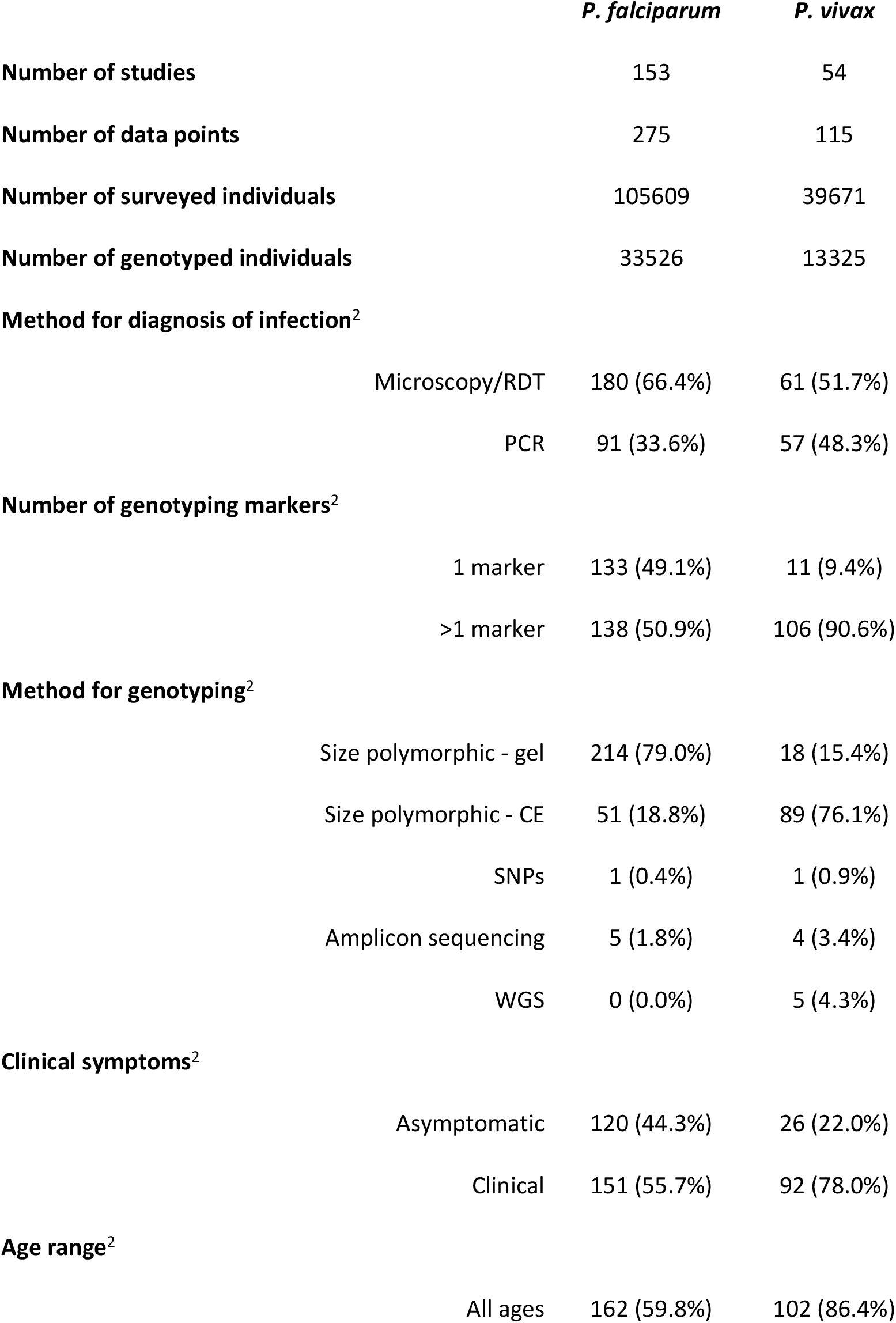
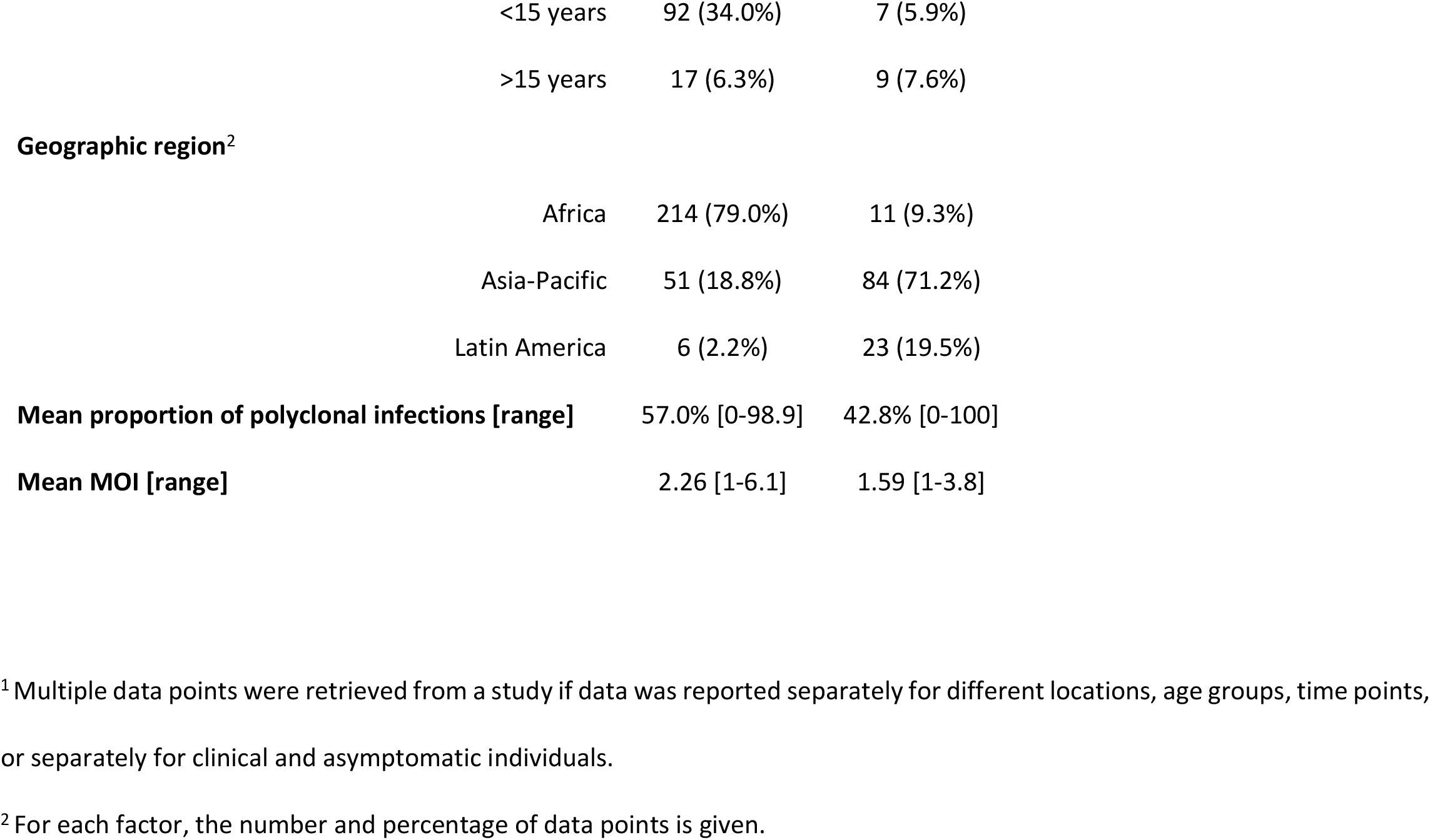
Characteristics of studies analyzed. **Table 1:** Summary statistics of studies analyzed

|  | *P. falciparum* | *P. vivax* |
| --- | --- | --- |
| **Number of studies** | 153 | 54 |
| **Number of data points** | 275 | 115 |
| **Number of surveyed individuals** | 105609 | 39671 |
| **Number of genotyped individuals** | 33526 | 13325 |
| **Method for diagnosis of infection<sup>2</sup>** |  |  |
| Microscopy/RDT | 180 (66.4%) | 61 (51.7%) |
| PCR | 91 (33.6%) | 57 (48.3%) |
| **Number of genotyping markers<sup>2</sup>** |  |  |
| 1 marker | 133 (49.1%) | 11 (9.4%) |
| >1 marker | 138 (50.9%) | 106 (90.6%) |
| **Method for genotyping<sup>2</sup>** |  |  |
| Size polymorphic - gel | 214 (79.0%) | 18 (15.4%) |
| Size polymorphic - CE | 51 (18.8%) | 89 (76.1%) |
| SNPs | 1 (0.4%) | 1 (0.9%) |
| Amplicon sequencing | 5 (1.8%) | 4 (3.4%) |
| WGS | 0 (0.0%) | 5 (4.3%) |
| **Clinical symptoms<sup>2</sup>** |  |  |
| Asymptomatic | 120 (44.3%) | 26 (22.0%) |
| Clinical | 151 (55.7%) | 92 (78.0%) |
| **Age range<sup>2</sup>** |  |  |
| All ages | 162 (59.8%) | 102 (86.4%) |

**Table 1:** Characteristics of studies analyzed.
|  |  |  |  |
| --- | --- | --- | --- |
|  | <15 years | 92 (34.0%) | 7 (5.9%) |
|  | >15 years | 17 (6.3%) | 9 (7.6%) |
| **Geographic region<sup>2</sup>** |  |  |  |
|  | Africa | 214 (79.0%) | 11 (9.3%) |
|  | Asia-Pacific | 51 (18.8%) | 84 (71.2%) |
|  | Latin America | 6 (2.2%) | 23 (19.5%) |
| **Mean proportion of polyclonal infections [range]** |  | 57.0% [0-98.9] | 42.8% [0-100] |
| **Mean MOI [range]** |  | 2.26 [1-6.1] | 1.59 [1-3.8] |

<sup>1</sup> Multiple data points were retrieved from a study if data was reported separately for different locations, age groups, time points, or separately for clinical and asymptomatic individuals.
<sup>2</sup> For each factor, the number and percentage of data points is given.

### *P. falciparum* proportion of polyclonal infections

In univariate analysis, a significant correlation between the proportion of polyclonal infections and prevalence was observed (Figure 1A). Each percentage point increase in prevalence resulted in an 0.34% increase in the proportion of polyclonal infections (n=174, *P*<0.001). At low prevalence, the proportion of infections carrying multiple clones ranged from very low to almost 100% (Figure 1A). At higher prevalence, very few studies found a low proportion of polyclonal infections. In only two out of 43 studies reporting prevalence rates >50% was the proportions of polyclonal infections <50%. Over the time period analyzed (1990-2019), a pronounced decrease in the number of clinical malaria cases was observed globally. Nevertheless, no significant impact of the year of study on the proportion of polyclonal infections was observed (n=174, *P*=0.869, Figure 2A).

**Figure 1:**
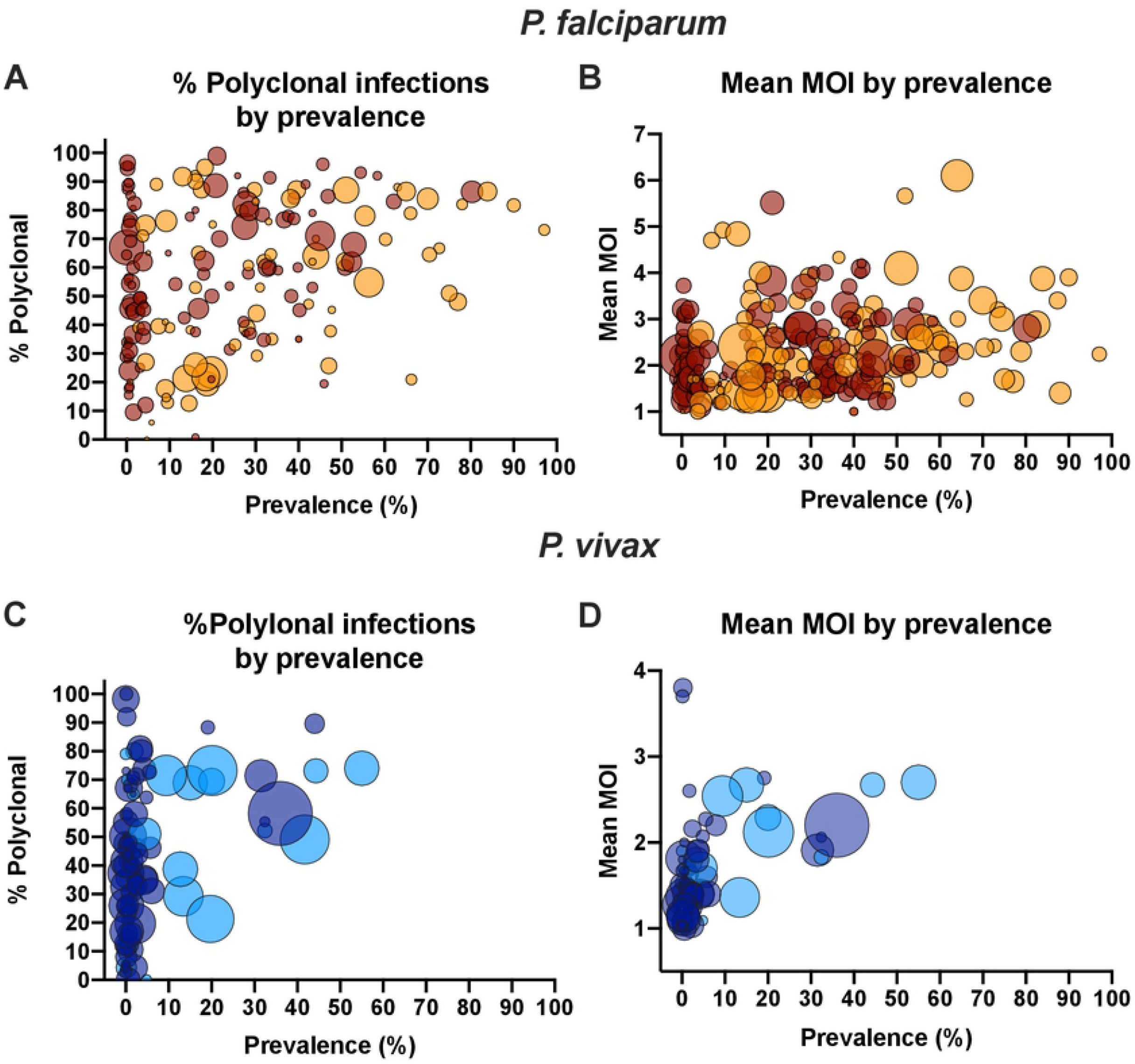
Relationship between prevalence and the proportion *P. falciparum* and *P. vivax* polyclonal infections (A, C) or mean multiplicity (B, D). Dot sizes represent the number of individuals genotyped. Dark red/dark blue = samples collected form asymptomatic individuals. Orange/light blue = samples collected form clinical patients.

**Figure 2:**
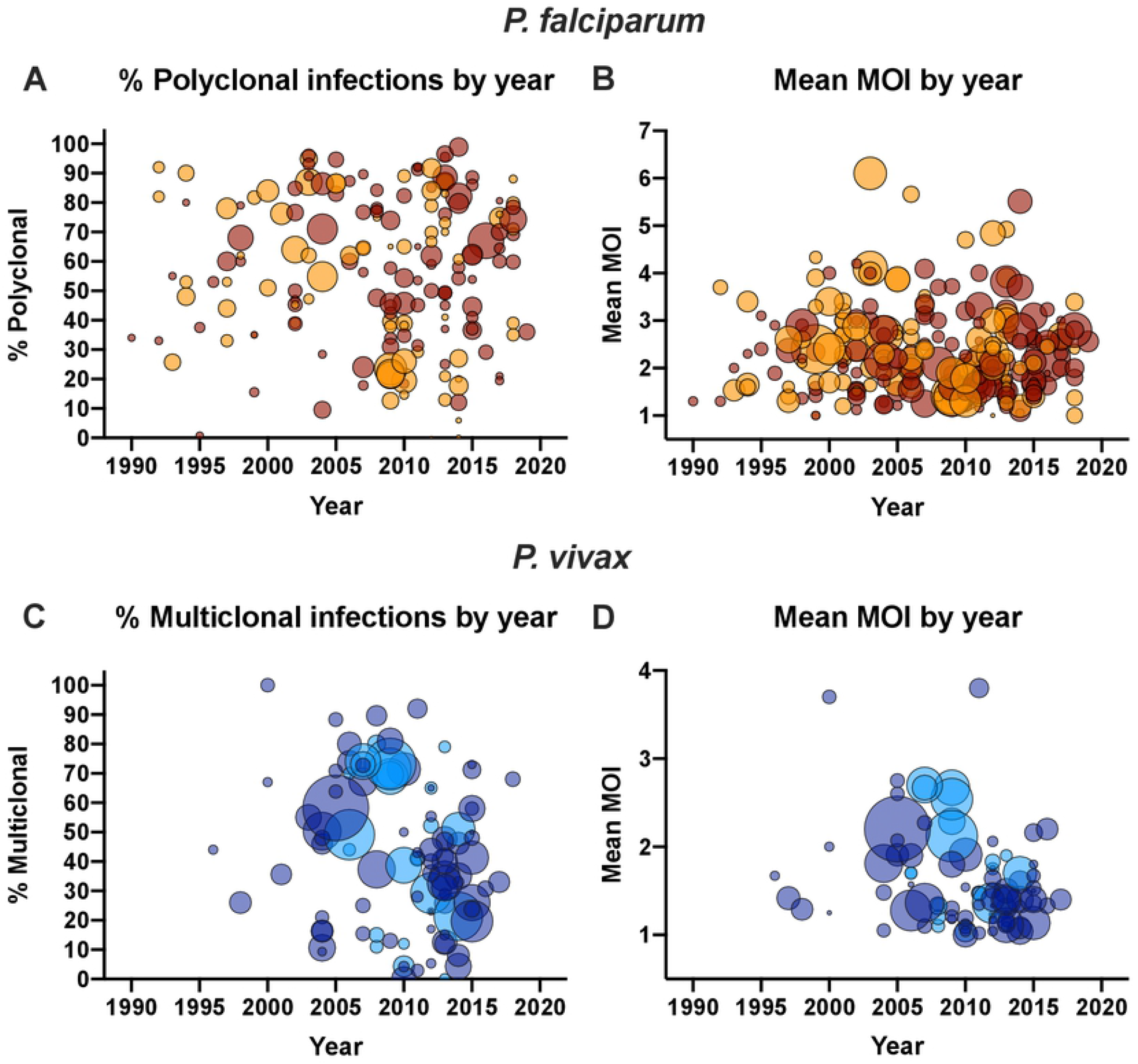
Relationship between year of sample collection and the proportion *P. falciparum* and *P. vivax* polyclonal infections (A, C) or mean multiplicity (B, D). Dot sizes represent the number of individuals genotyped. Dark red/dark blue = samples collected form asymptomatic individuals. Orange/light blue = samples collected form clinical patients.

The proportion of infections diagnosed by PCR that were polyclonal was 10.4% less than for infections diagnosed by microscopy or RDT (irrespective of whether individuals showed symptoms) (n=174, *P*=0.010, Figure 3A). Typing multiple markers resulted in 8.2% more polyclonal infections (n=174, *P*=0.032, Figure 3B). When size-polymorphic markers were typed, typing by CE instead of on gel resulted in detection of 12.8% fewer polyclonal infections (n=168, *P*=0.007, Figure 3C). Notably, the proportion of studies that used CE was 3-fold higher in the Asia-Pacific region compared to Africa (42.9 vs. 13.2%, *P*<0.001). In the Asia-Pacific region, the proportion of polyclonal infections was 21.9% lower than the proportion in Africa (n=168, *P*<0.001, Figure 3F).

**Figure 3:**
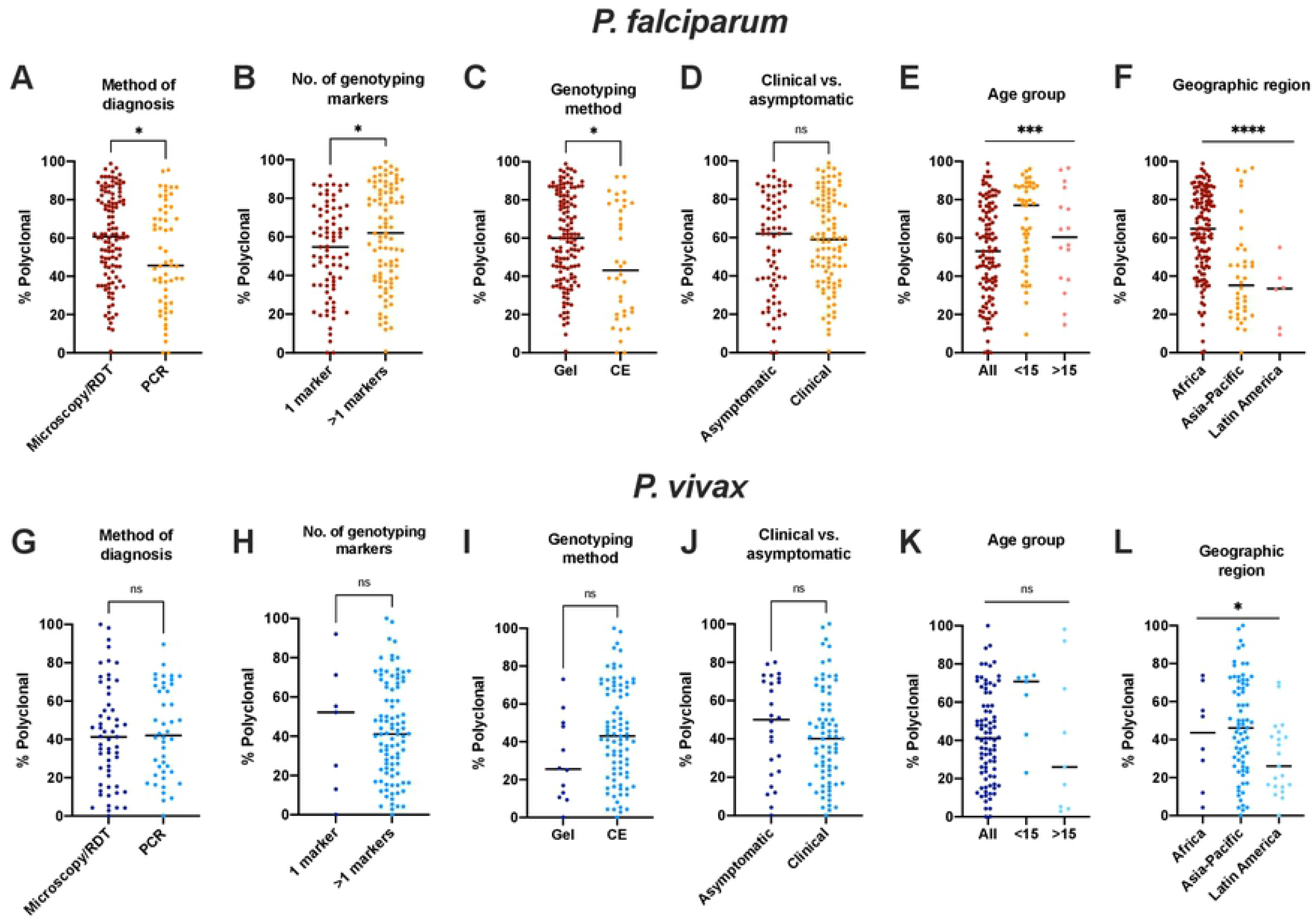
Impact of presence of laboratory methods (method for diagnosis, number of genotyping markers, method to size size-polymorphic amplicons) and patient characteristics (presence of clinical symptoms, age group, geographic region) on the proportion *P. falciparum* (A-F) and *P. vivax* (G-L) polyclonal infections. Black line denotes median. Groups were compared by Mann-Whitney test (if 2 groups), or regression analysis (if 3 groups). * *P* ≤ 0.05, \*\**P* ≤ 0.01, *** *P* ≤ 0.001, *** *P* ≤ 0.0001, ns = not significant.

No significant difference was observed in the proportion of polyclonal infections between samples collected from clinical and asymptomatic individuals (n=174, *P*=0.358, Figure 3D). Age group had a significant impact on the proportion of polyclonal infections, with the highest proportion observed in studies only enrolling children (n=174, *P*=0.0033, Figure 3E).

In multivariable analysis, the year the study was conducted, sizing by CE vs. gel, age group, and whether samples were from clinical or asymptomatic individuals did not reach significance and were excluded from the final model (Table 2A). In the reduced model, each percentage point increase in prevalence resulted in an 0.28% increase in the proportion of polyclonal infections (*P*=0.001). Typing multiple markers instead of only one marker resulted in detection of 14.1% more polyclonal infections (*P*<0.001). Diagnosis of infections by PCR instead of microscopy or RDT resulted in 9.4% fewer polyclonal infections detected (*P*=0.013). Compared to Africa, the proportion of polyclonal infections in the Asia-Pacific region was 17.8% lower, and in Latin America, it was 28.0% lower (*P*<0.001).

**Table 2:**
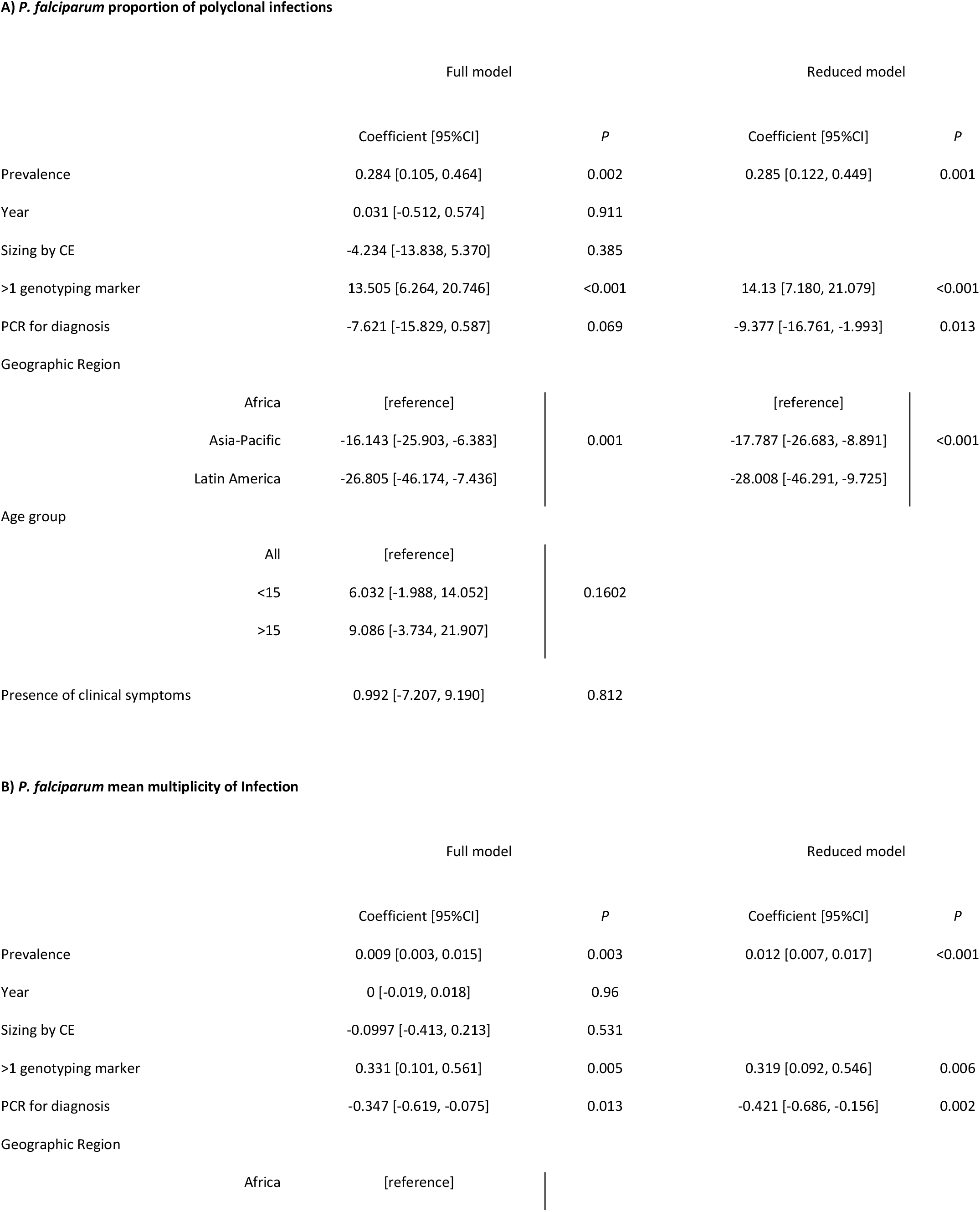
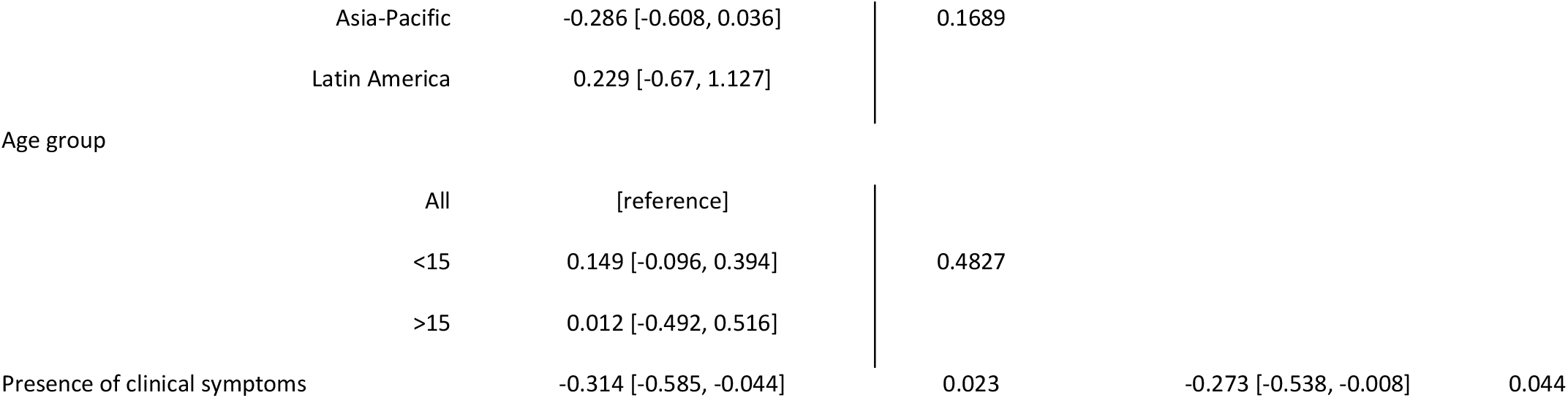
Multivariable predictors of A) *P. falciparum* proportion of polyclonal infections, and B) mean multiple clone infections. 95%CI = 95% confidence interval.

### *P. falciparum* mean multiplicity

In univariate analysis, each percentage point increase in prevalence resulted in an increase in mean MOI of 0.010 (n=254, *P*<0.001, Figure 1B). No impact of the year of study on mean MOI was observed (n=254, *P*=0.241, Figure 2B).

Mean MOI did not differ significantly between studies on clinical and asymptomatic individuals (n=254, *P*=0.158). A trend towards lower mean MOI was observed in studies using PCR for diagnosis compared to microscopy or RDT (mean MOI 0.219 lower if PCR used, n=254, *P*=0.069). In studies including children <15 years (n=90), mean MOI was 0.350 higher compared to studies including all ages (n=150, *P*=0.004). No difference was observed between studies including only adults >15 years (n=14) and all ages (*P*=0.988). Typing of size-polymorphic markers by CE instead of gel resulted in a trend toward lower mean MOI (0.270 lower by CE, n=249, *P*=0.073).

In multivariable analysis, higher prevalence and typing multiple markers resulted in higher mean MOI. Diagnosis by PCR and typing clinical samples resulted in lower mean MOI (Table 2B).

### *P. falciparum* marker *msp2*

While a wide variety of genotyping markers was used for *P. falciparum* genotyping, *msp2* was the most frequently used marker. 97 data sets reported the proportion of polyclonal infections based on *msp2*, and 172 data sets reported mean multiplicity based on *msp2*. To study whether typing the same marker impacts the relationship between multiplicity and prevalence, the analysis was repeated using *msp2* only.

The proportion of polyclonal infections increased by 0.32% per percentage point increase in prevalence (n=97, *P*=0.002, Figure 4A). As for the entire dataset, at low prevalence, the proportion of polyclonal infections varied from 0 to almost 100%. At high prevalence, in most studies, the proportion was >50%. Mean MOI based on *msp2* increased by 0.012 per percentage point increase in prevalence (n=172, *P*<0.001, Figure 4B).

**Figure 4:**
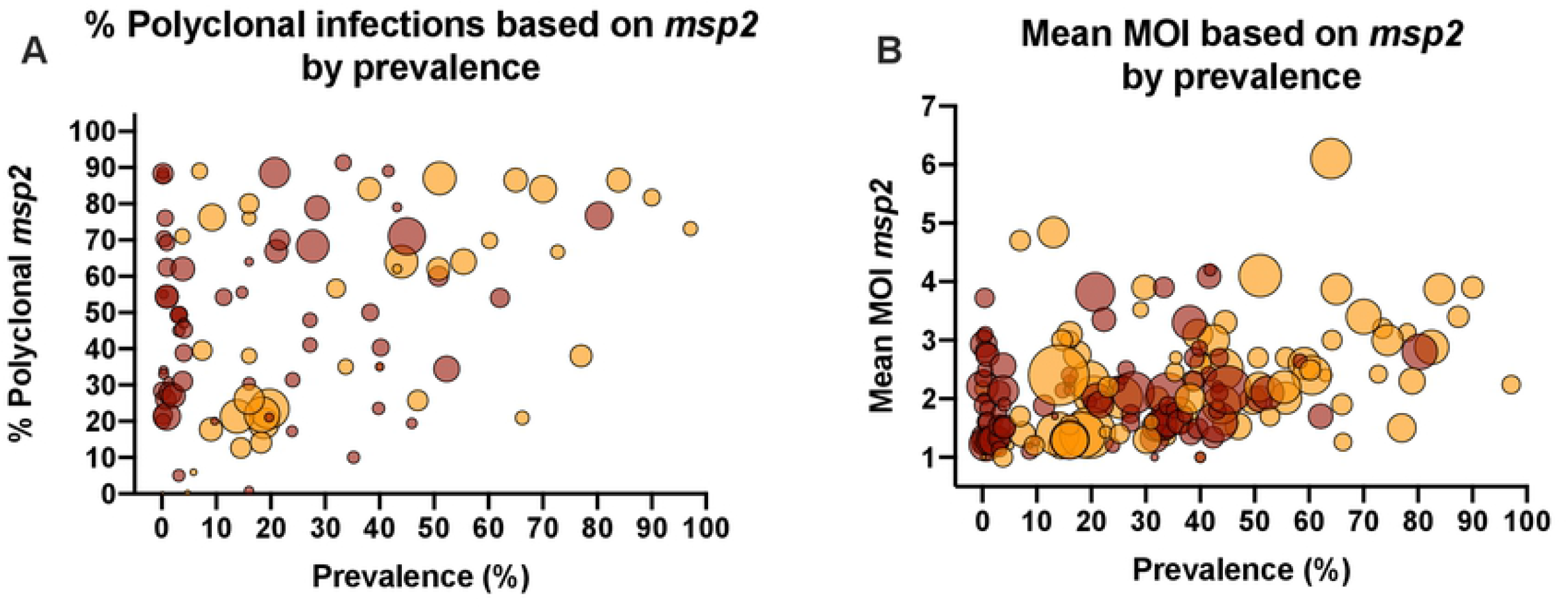
Relationship between the proportion *P. falciparum* polyclonal infections (A) or mean multiplicity (B) based on typing marker *msp2* and prevalence. Dot sizes represent the number of individuals genotyped. Dark red = samples collected form asymptomatic individuals. Orange = samples collected form clinical patients.

### *P. vivax* literature search

Using a broad set of search criteria, 54 studies were identified that met inclusion criteria, yielding 115 data points that included 13,325 genotyped individuals (Table 1). On average 118 individuals were genotyped per study. The proportion of polyclonal infections ranged from 0-100%, and the mean multiplicity ranged from 1-3.8.

### *P. vivax* proportion of polyclonal infections

As for *P. falciparum*, at low *P. vivax* prevalence, the proportion of polyclonal infections varied widely from almost 0% to close to 100% (Figure 1C). Few studies were available where *P. vivax* prevalence was high, and the proportion of polyclonal infections was high in all of these studies. In univariate analysis, each percentage point increase in prevalence resulted in an 0.78% increase in the proportion of polyclonal infections (n=101, *P*<0.001). Over the time period analyzed (1996-2018), a trend toward a lower proportion of polyclonal infections was observed (1.02% reduction per year, n=99, *P*=0.070, Figure 1C).

In univariate analysis, no difference was observed in the proportion of polyclonal infections whether diagnosis was by microscopy/RDT or PCR (n=101, *P*=0.764, Figure 3G), using multiple genotyping markers (n=10, *P*=0.917, Figure 3H), typing by CE vs. gel (n=90, *P*=0.082, Figure 3J), between asymptomatic and clinical infections (n=101, *P*=0.418, Figure 3J), or among age groups (n=101, *P*=0.169, Figure 3K). A significant difference was observed among geographic regions (n=101, *P*=0.028, Figure 3L).

In multivariable analysis, each percentage point increase in prevalence resulted in 0.48% more polyclonal infections (*P*<0.001, Table 3A). Compared to Africa, in the Asia-Pacific the proportion of polyclonal infections was 9.6% higher, and in Latin America 7.5% lower (*P*=0.0141, Table 3A). Few studies typed a single marker, which resulted in 35% fewer polyclonal infections detected (*P*=0.006, Table 3A). Further significant predictors were year of sample collection and age group (Table 3A).

**Table 3:**
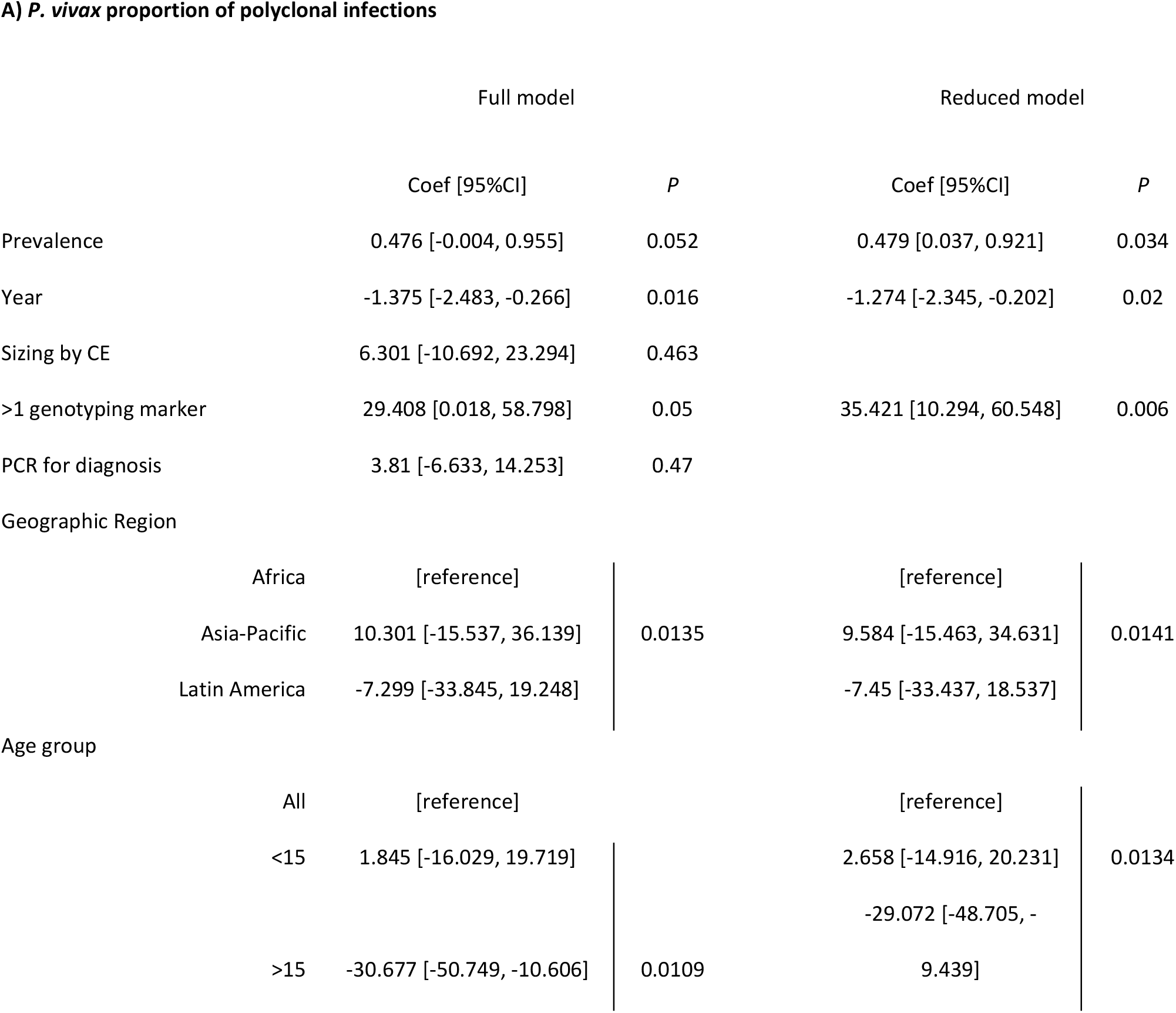
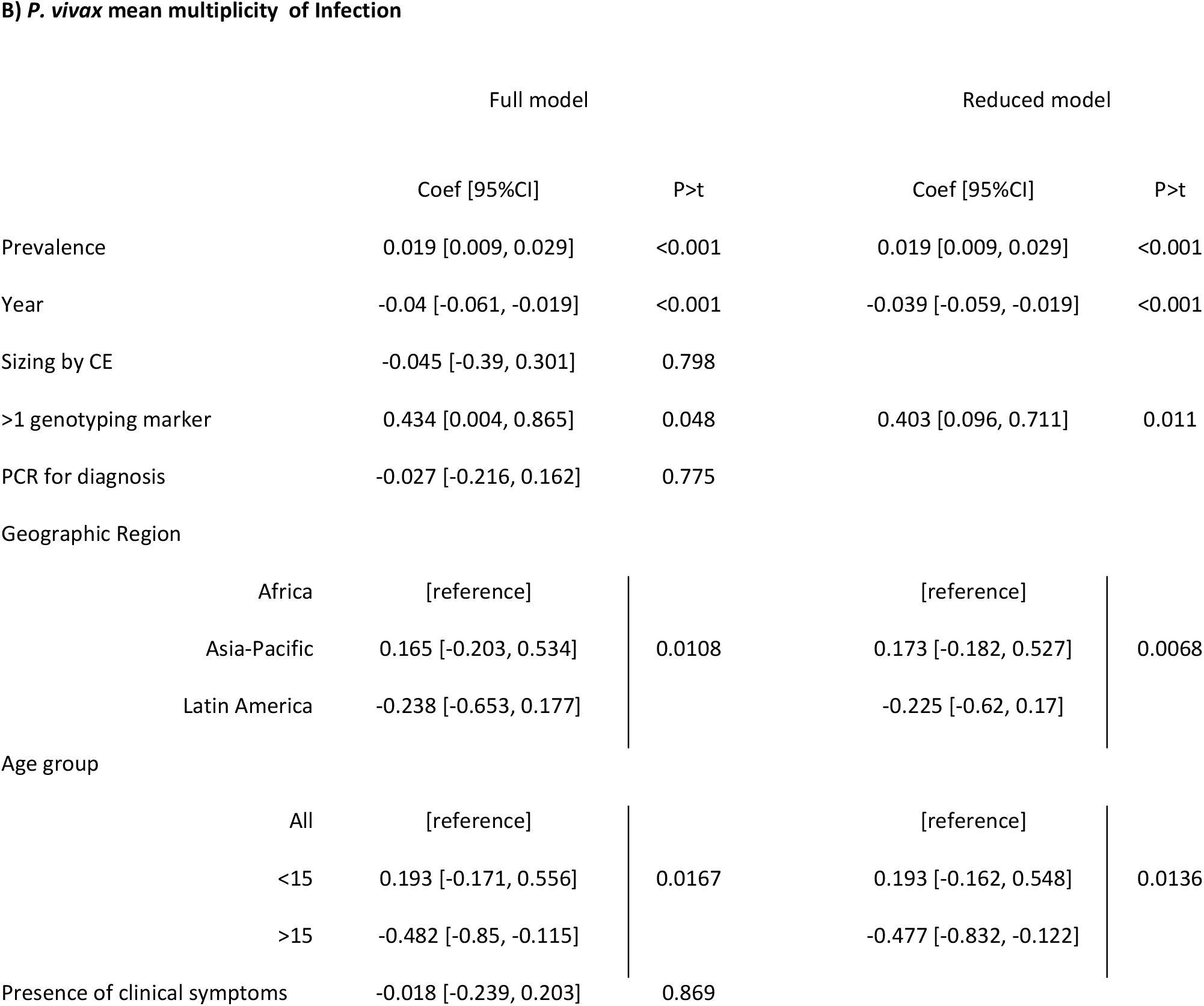
Multivariable predictors of A) *P. vivax* proportion of polyclonal infections, and B) mean multiple clone infections. 95%CI = 95% confidence interval.

### *P. vivax* mean multiplicity

In univariate analysis, each percentage point increase in prevalence resulted in an increase in mean MOI of 0.025 (n=91, *P*<0.001, Figure 1D). The year of study was not a significant predictor for mean MOI (n=91, *P*=0.207, Figure 2D). Presence of clinical symptoms (n=91, *P*=0.169) and method of diagnosis (n=91, *P*=0.999) did not impact mean MOI. Mean MOI was higher in studies enrolling only children (n=77, mean MOI=2.26) compared to all ages (n=6, mean MOI=1.54, *P*=0.002). In multivariable analysis, the same predictors as for the proportion of polyclonal infections were significant, namely prevalence, year of sample collection, typing multiple markers, geographic region, and age group (Table 3B).

## Discussion

MOI and the proportion of polyclonal infections are among the most frequently reported measures of malaria genomic epidemiology studies and have been proposed as surrogate markers for transmission intensity. Analysis of data from a large number of studies showed that polyclonal infections were frequent in almost all studied populations, even in low transmission settings. A weak correlation between multiplicity indices and prevalence was observed. For both species, once prevalence reached 40%, the majority of infections were polyclonal in almost all studies. At lower prevalence, some studies found a low proportion of polyclonal infections, while in others it was very high.

Multiplicity indices were lower in *P. falciparum* infections diagnosed by PCR instead of microscopy or RDT. Detectability of clones decreases as parasite densities decrease [30]. While parasite densities were not reported by most studies and thus not analyzed, the lower multiplicity when PCR was used for diagnosis can likely be explained by lower mean parasite densities in these infections. However, despite generally higher mean parasite densities in clinical cases compared to asymptomatic individuals, no difference in the proportion of polyclonal infections was observed between the two groups. Asymptomatic infections can persist for many months. Superinfections during this time can result in high multiplicity [3, 34]. A previous study comparing asymptomatic and clinical infections in West Papua from the same site indeed found higher multiplicity in asymptomatic individuals [24]. Possibly, in the data set analyzed here, higher detectability of clones in high-density clinical infections was balanced by higher multiplicity in asymptomatic individuals, resulting in overall similar estimates of the proportion of polyclonal infections in clinical and asymptomatic infections.

Estimates of multiplicity are heavily impacted by the ability to detect minority clones and by the discrimination power of the genotyping method selected. Size polymorphic markers were the preferred method for both species, but pronounced differences in the methods used for *P. falciparum* and *P. vivax* were observed. For *P. falciparum*, sizing of polymorphic markers on gel remains the primary method for genotyping. 79% of all studies sized PCR amplicons on gel, with little change in more recent studies (74% in studies conducted since 2015). For *P. vivax*, 76% of studies used CE for sizing. The choice of gel vs. CE can affect multiplicity estimates for two reasons. First, capillary electrophoresis increases detectability of minority clones. Second, it allows distinction of fragments of similar size compared to visualization on gel. By CE, fragments differing as little as 3 base pairs in size can be distinguished. In contrast, when binning using gels, resolution might be as low as 50 base pairs [35]. Previous studies comparing the two methods directly showed that CE yields higher multiplicity [31, 32]. In the current study, however, in univariate analysis typing by CE resulted in a lower proportion of *P. falciparum* polyclonal infections, and the method was not a significant predictor in multivariable analysis. The result from the univariate analysis could be explained by the more widespread use of CE in the Asia-Pacific compared to Africa, where transmission and multiplicity are generally lower than in Africa.

For *P. falciparum* genotyping, 49% of studies typed a single marker. In multivariable analysis, typing multiple *P. falciparum* markers increased the proportion of polyclonal infections detected by 14%. An even stronger increase of 35% was observed for *P. vivax*, where only 9% of studies typed a single marker.

Pronounced differences were observed among geographic regions. For *P. falciparum*, the proportion of polyclonal infections was highest in Africa, and for *P. vivax*, in the Asia-Pacific. In both cases, this mirrors high transmission intensity.

Marker *msp2* has been widely used for *P. falciparum* genotyping, allowing to repeat the analysis on this marker only. Results mirrored those of the entire dataset. The relationship between the proportion of polyclonal infections and prevalence remained poor. Despite its wide use, inclusion criteria for minority clones when typing *msp2* differ among laboratories, and are often not reported. When using CE, the criteria applied include an arbitrary cut off of 33% of the height of the main peak [6], a cut off based on the signal intensity of the size standard [36], or inclusion of all peaks above a set fluorescent signal threshold [17]. This variation in criteria to include minority clones likely adds to the weak relationship between multiplicity estimates and transmission intensity.

In many studies, in particular those typing samples from clinical patients, no data on transmission intensity was given, such as population prevalence, clinical incidence, or test positive rate. In order to conduct more detailed analysis of the relationship between multiplicity and transmission intensity, it is recommended to report such measures along with genotyping data.

In conclusion, the weak correlation between mean MOI or the proportion of polyclonal infections and prevalence indicates these measures do not accurately stratify transmission intensity when different studies are combined. Further studies will be required to understand the factors affecting the presence of polyclonal infections when prevalence is low. The possibility of a high proportion of polyclonal infections even in low transmission sites should be considered when typing markers of drug resistance or *hrp2* deletion. In polyclonal infections the presence of deletions can be masked by wild-type infections [21], and the frequency of mutations conferring drug resistance observed by genotyping might not accurately reflect the prevalence among all clones [19].

## Supporting Information

Supplementary File S1: Data analyzed for this study.

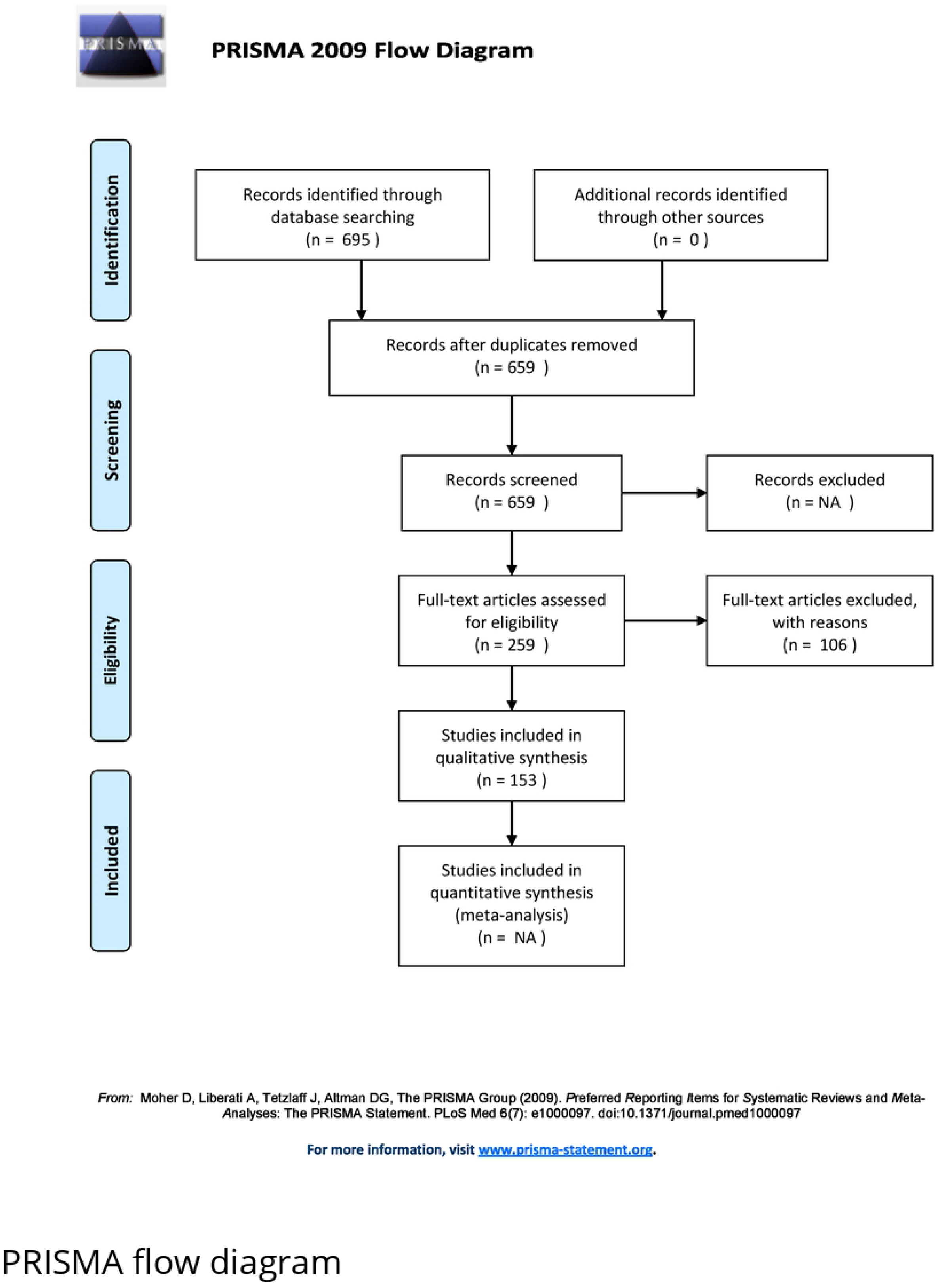

